# GENET: AI-Powered Interactive Visualization Workflows to Explore Biomedical Entity Networks

**DOI:** 10.64898/2025.12.12.694029

**Authors:** Bum Chul Kwon, Natasha Mulligan, Joao Bettencourt-Silva, Ta-Hsin Li, Bharath Dandala, Feng Lin, Ching-Huei Tsou, Pablo Meyer

## Abstract

Formulating hypotheses about gene-disease associations requires logical inference from prior data, followed by a laborious literature review. AI models trained on curated datasets (e.g., GWAS Catalog) can suggest SNP–disease links, but validating these predictions still demands manual evidence extraction. To streamline this process, we present GENET (Genomic Evidence Network Exploration Tool), an AI-enhanced, end-to-end visual analytics workflow applied to Age-Related Macular Degeneration (AMD). GENET comprises four sequential steps: (1) biomedical network analysis: a dual-encoder neural model identifies genes or SNPs associated with a target disease and vice versa; (2) literature evidence mining pipeline: a pipeline retrieves relevant papers and, using large language models, extracts biomedical entities and relations; (3) clustering: embeddings from pre-trained biomedical language models (BioBERT, BioLinkBERT) are clustered to group related concepts; (4) interactive visualizations: clusters and their networks are visualized with interactive features for hypothesis testing and insight generation. The workflow enables iterative hypothesis formulation and evidence validation, uncovering novel associations. GENET is open-sourced (https://github.com/BiomedSciAI/genet) and available for demonstration at https://genet.pythonanywhere.com.

## Introduction

The rapid expansion of homogeneously annotated knowledge of the genetic causes of diseases from Genome Wide Association Studies (GWAS) and the ever-growing corpus of biomedical literature have created unprecedented opportunities for understanding genetic determinants of complex diseases. Single-Nucleotide Polymorphisms (SNPs) identified in large-scale association catalogs such as the GWAS Catalog and ClinVar (Sollis et al., 2023; Landrum et al., 2014) are routinely mined to build models and generate hypotheses about disease-associated genes (Piñero et al., 2016), leading to disease mechanisms (Frazer et al., 2021), novel drug targets, and patient stratification strategies. In practice, however, moving from a statistical signal or a new gene-disease predicted association to a biologically plausible hypothesis remains a labor-intensive endeavor. Researchers must first analyze the inundating, raw association data, then manually retrieve, read, and synthesize evidence from a plethora of publications and expert-curated resources to assess the credibility of a gene-disease link. This iterative cycle of association analysis, literature mining, and knowledge integration and exploration is a major bottleneck that limits the speed at which novel genotype-phenotype relationships can be discovered, validated, and translated into therapeutic insights. Conversely, it is also challenging for domain experts to write code for configuring various computational pipelines with this goal in mind.

Furthermore, machine learning models trained on sparse, known disease labels have been considered insufficiently reliable to predict plausible new gene-disease associations that were not captured in the original GWAS analyses (Frazer et al., 2021). However, recent advances in artificial intelligence (AI) have demonstrated that biomedical foundation models pretrained with a large scale of DNA sequences can be used to predict new gene-disease associations (Frazer et al., 2021; Li et al., 2025). In addition, large language models (LLMs) adapted for biomedical scientific literature, such as BioBERT and BioLinkBERT (Lee et al., 2020; Yasunaga et al., 2022), can further enable the automated extraction of biomedical entities and their relational statements from unstructured text. While these technologies can surface candidate associations and retrieve relevant literature at scale, they do not yet provide a cohesive, user-driven environment in which scientists can explore, validate, and refine the new hypotheses generated by these new types of models through visual interaction with the underlying evidence. Existing pipelines either focus on isolated tasks—such as text mining or network inference—without offering an integrated workflow that bridges predictive modeling, evidence collection, and exploratory analysis.

To address these challenges, we present GENET (Genomic Evidence Network Exploration Tool), an AI-powered, end-to-end visual analytics workflow designed explicitly for exploring biomedical entity networks in scientific literature and databases. GENET orchestrates four tightly coupled, sequential steps, where each feeding into the next: (i) Biomedical network analysis, where a network-based inference engine identifies genes and SNPs that are indirectly linked to a disease of interest through shared network neighborhoods; we call it SNP2Trait (ii) Literature evidence mining, which automatically retrieves pertinent publications and applies LLM-driven entity-relation extraction to construct a structured knowledge graph; (iii) Embedding-based clustering, leveraging pre-trained biomedical language models to embed extracted entities and relations, followed by unsupervised clustering that reveals coherent thematic groups; and (iv) Interactive visualizations, offering coordinated views of entity clusters, network topology, and supporting evidence, together with interactive handles for filtering, drilling down, and iteratively refining hypotheses. Each stage consumes the output of the preceding step, enabling a seamless feedback loop between computational inference and human expertise.

By integrating AI foundation models, automated, LLM-based literature mining, and interactive visualizations, GENET empowers researchers to rapidly formulate, evaluate, and iterate on SNP-disease hypotheses against a rich backdrop of curated evidence. In the remainder of this paper, we detail the workflows, describe individual components, and demonstrate the utility of GENET through a use case centered around Age-Related Macular Degeneration (AMD). GENET shows how a tightly coupled human-AI workflow can enable the discovery of biologically meaningful genetic associations supported by evidence in biomedical literature.

## Materials and Methods

GENET is an AI-enhanced visual analytics workflow that consists of four distinct and connected steps that help users to first discover new putative SNP-disease associations, then collect evidence from scientific literature supporting the predicted associations, extract knowledge from it as biomedical entity networks, and finally interactively explore them using visualizations. GENET is released as open-source software (see https://github.com/BiomedSciAI/genet). A web deployment of the visualization component is also provided for demonstration purposes (see https://genet.pythonanywhere.com/). In this section, we provide an overview of the four steps.

### Overall Workflow

This workflow allows domain experts to explore biomedical entity networks through interactive, evidence-driven visual analytics systems. Figure 1 shows an overview of the workflow that consists of four key steps, each building on the previous step. The computational pipeline begins with SNP2trait, the biological network analysis identifying key genes, SNPs, or traits, including diseases that are potentially associated with given entities. Then, the Literature Evidence Mining Pipeline extracts and annotates relevant entity relationships from public databases and scientific literature. In the following step, we use pre-trained biomedical language models to generate embeddings for the extracted entities and cluster them to reveal structural patterns. Finally, GENET visualization allows users to visually navigate and interrogate these networks, facilitating hypothesis generation and insight discovery.

**Figure 1.**
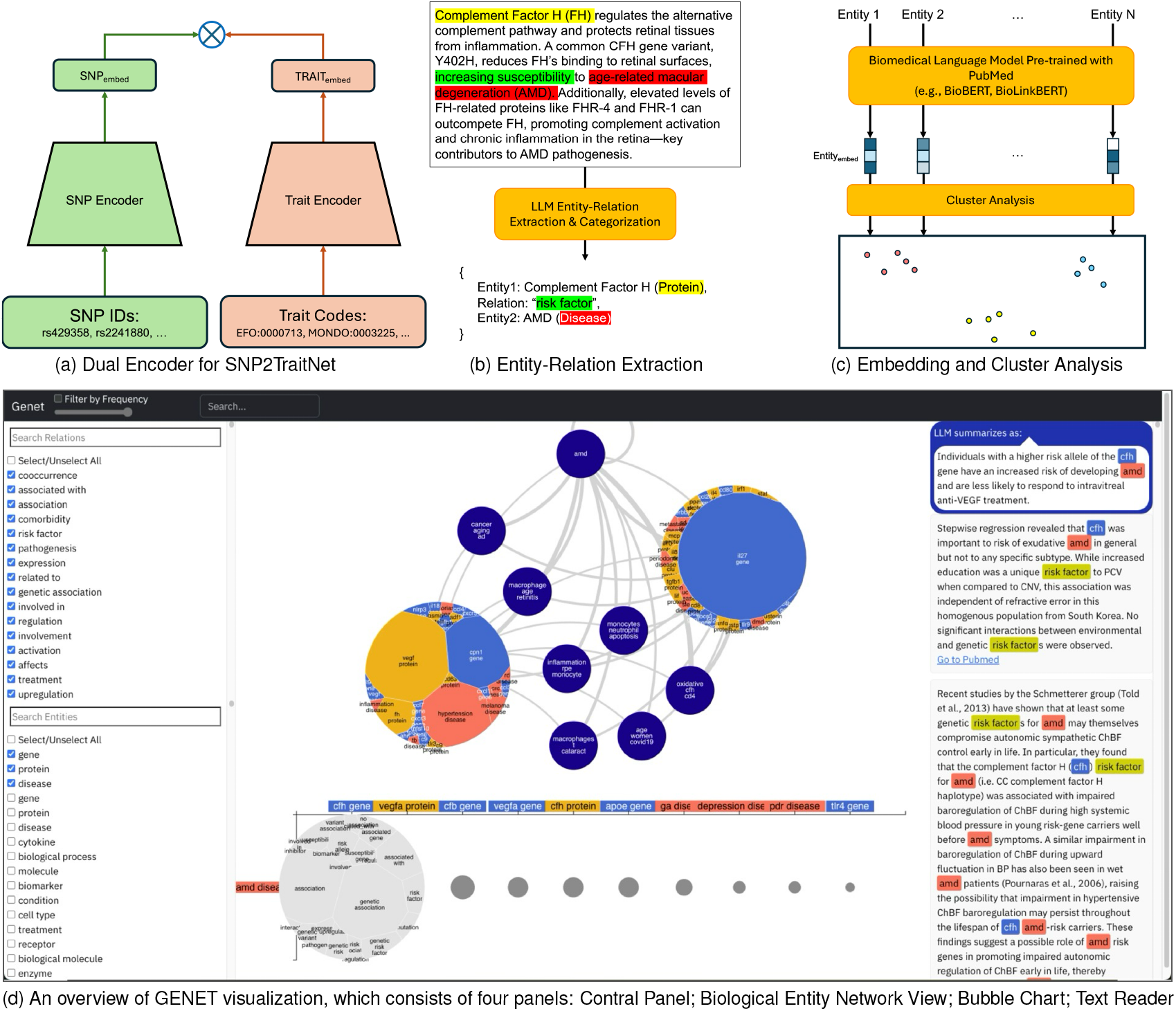
GENET workflow overview: four key steps: (a) Biological Network Analysis using SNP2TraitNet: identify genes/SNPs associated with traits or vice versa; Literature Mining Pipeline: extract entities and their relations from scientific literature; (c) Embedding Generation and Cluster Analysis: generate embeddings using pre-trained biomedical lanaguage models and cluster entities to discover semantic groups; (d) GENET Visualization: visually explore biomedical entity networks.

### Identify associations between SNPs and diseases

The first step is to predict links between target diseases and genes/SNPs (see Figure 1(a)). To achieve it, we adapted the dual-encoder model architecture (Ni et al., 2022) to estimate the probability of an SNP being associated with a given disease: P(*S* | *D*) as well as that of a disease being associated with a given SNP: P(*D* | *S*). We used ClinVar (Landrum et al., 2019) and GWAS Catalog (Landrum et al., 2014), which contain public archives of genome-wide association studies between DNA sequence variants (i.e., a SNP) and traits that include diseases, to train a shallow neural network model with a hidden layer using the SGNS algorithm. This is in itself innovative as in general, the well-established problem of SNP-to-disease association is typically approached using statistical methods. In total, we collected 1,537,516 pairs of SNPs and Traits, where 548,627 unique SNPs in the form of the Reference SNP Cluster ID (i.e., RS ID) and 16,014 Trait codes in 12 different ontologies were present, namely: EFO, GO, HP, MONDO, Orphanet, HANCESTRO, NCIT, OBA, PATO, MedGen, OMIM, MeSH. Two different encoders with two-layered neural networks were trained for SNPs and Traits, respectively, and simultaneously, where each network generates embeddings with the same dimension for every SNP and Trait in the latent space so that cross-entropy loss on SNP-Trait pairs is minimized. Since our dataset only contains positive associations, we randomly chose 10 negative samples for each SNP. We chose the hyperparameter setting that yielded the best performance on the validation set while keeping the test set untouched. The following statistics summarize the performance of our model on the test dataset: 1) accuracy: .96; 2) AUCROC: .97; 3) Precision: .79; 4) Recall: .73. To use the trained model in the workflow, we implemented a script that identifies the associations between SNPs and Traits in three different ways: i) SNP to Traits; ii) Trait to SNP/Gene; iii) Gene Name to Trait. Since our model was trained with RS IDs, we implemented a script to translate the RS IDs to gene names and vice versa. In the inference time, we use the trained model to generate embeddings for SNPs for a given target trait/disease, and then calculate the cosine distance from the SNPs to other known SNPs associated with the target trait. In order to predict new SNP to Trait putative associations, we chose the SNPs that are closest to the given trait in the shared latent space and vice versa.

### Extract entities and relations from literature

Inspired by our previous literature evidence mining pipeline (Bettencourt-Silva et al., 2024), we implemented a fast workflow to collect abstracts relevant to the set of genes, diseases, and SNPs identified in the previous step from scientific literature and to extract and label biomedical entities and relations from each article (see Figure 1(b)). After finding potentially associated genes, diseases, or SNPs in the previous step, independent searches were performed. We implemented a script that fetches articles relevant to users’ queries by using PubMed Entrez Programming Utilities (E-utilities) (Sayers, 2022). Then, we used an LLM (Granite-4-Tiny (IBM Research, 2025)) with the Mellea (Mellea Contributors, 2025) framework to extract the triplets of entity1, relation, and entity2 from each abstract of collected articles. For the LLM call, we set the following requirements in the form of a prompt: 1) all extracted entities must be biomedical or scientific terms, expressed as nouns or noun phrases; 2) all relations must be in simplified, canonical form; 3) all extracted entities and relations need to be strictly derived from the given abstract without using any external knowledge; 4) output must be in the JSON format. Within the five maximum attempts for each abstract, we validated the results using LLM-as-a-Judge and re-invoked the LLM with the same prompt. The resulting JSON list of triplets was then fed to the categorization step, where LLM was given examples to label entities: e.g., Entity: BRCA1 → gene, Entity: ApoE4 → allele, Entity: metformin → drug/pharmaceutical. In addition, the source information of the extracted knowledge, such as document link and text chunks where the entities appear together, is also stored.

### Embedding generation and cluster analysis

With collected lists of entities and relations extracted from biomedical literature, we perform clustering to group them (see Figure 1(c)). We employ pre-trained biomedical language models like BioBERT (Lee et al., 2020) or BioLinkBERT (Yasunaga et al., 2022) to generate semantically meaningful embeddings for entities and relations. Then, we run clustering on the list of entities and relations separately using the K-Means algorithm and a user-chosen parameter for the number of clusters. We annotate the JSON list of entities and relations with the clustering information.

### Visual exploration of biomedical entity networks

With the evidence data file enriched with clustering results in the previous step, GENET provides the control panel (Figure 1 (d), Left) and the biomedical entity network view (Figure 1 (d), Top Center). The control panel provides two lists of checkboxes, containing unique entities and relations, respectively. Checking or unchecking the items will set corresponding filters to the underlying evidence datasets to be visualized so that users can focus on a set of interested biomedical entities and relations at a time. In the navigation bar at the top, users can adjust the filter slider to randomly sample data by user-defined proportions and search for entities and relations by their names.

The main view provides a node-link diagram (Figure 1 (d), Top Center), where each node is a cluster of entities, and each edge represents a link between a pair of clusters. Here, the edge between clusters is the accumulation of all pairwise relations between entities from Cluster A and those from Cluster B. The edge width is proportionally corresponding to the number of relations. Hovering over each edge shows a ranked list of the top N most frequently appearing relations. Each node initially provides labels with up to three most frequently occurring entities. Users can hover over each cluster to view the expanded list of the top N most frequently appearing entities in the cluster. Clicking on a node expands it to a Voronoi treemap (Balzer and Deussen, 2005), where the size of each entity polygon represents the relative frequency. Across the views, GENET uses a consistent color map for entity types, e.g., gene, protein, disease.

Users can either click on the edge between clusters or an entity polygon within the Voronoi treemap to open the bubble plot at the bottom as Figure 1 (d), Bottom Center shows. It shows a list of seed entities, used in the evidence pipeline as rows, and connected entities as columns. The bubbles correspond to the number of relations between the entities in the corresponding row and column. Clicking on each bubble opens a Voronoi treemap, this time showing relation polygons within. By viewing this, the users can inspect the relation types between a pair of entities.

Clicking on one relation type between a pair of entities opens the text reader view on the right side (Figure 1 (d), Right). The text reader view opens all excerpts, the evidence that includes the particular relations users clicked. The text reader also highlights the entities and relations with corresponding colors. In the background, to help users quickly understand the gist of the excerpts, we use an LLM (e.g., Granite-4.0-Tiny) to summarize the excerpts with respect to the entities and relations, and the results are appended to the list of excerpts on top. Each excerpt provides a link to the full text in PubMed so that users can find and read the paper easily.

The text reader provides additional, entity-specific visualizations for SNPs, proteins, and molecules. Users click on the entities to retrieve source data using external databases or services and then invoke corresponding visualizations at the center. In the case of SNPs, we use the Entrez programming utilities, i.e., eutils, provided by the National Library of Medicine (Sayers, 2010) to fetch chromosome positions of the variant in the human genome. Then, we use IGV.js (Robinson et al., 2023) to visualize the genomics viewer (Figure 2 (a)). When users click on protein entities, GENET fetches the PDB file, if available, by the provided names using RCSB (Berman et al., 2000) and then visualizes it using the NGL Viewer (Rose and Hildebrand, 2015) embedded at the center of the screen (Figure 2 (b)). For molecule entities, we look up the SMIlES representation of target molecules by using PubChem (Kim et al., 2016) and visualize them using RDKit (Bento et al., 2020) (Figure 2 (c)). These individual, entity-specific viewers help users further explore the structure and function of proteins and molecules, and genomic data visually.

**Figure 2.**
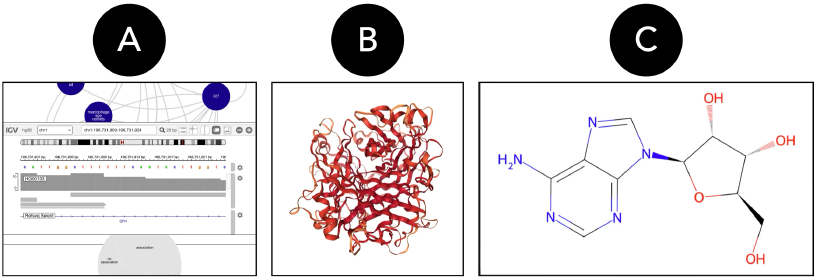
In the text reader, users can click on entities to open the following views: (a) Genome Browser using IGV.js (Robinson et al., 2023); (b) Protein Visualization using NGL Viewer (Rose and Hildebrand, 2015); (c) Molecule Visualization using RDKit (Bento et al., 2020).

### Use Case

In this use case, we applied GENET to the important disease of age-related macular degeneration (AMD). AMD causes the loss of a person’s central vision, affecting an estimated older population of 200 million worldwide. Several genes have been associated with its faster progress (Fritsche et al., 2016), in particular, genes involved in the complement system, an important part of the innate immunity, such as *CFH* (Klein et al., 2005) *C5* and *C3*, key inflammatory proteins activated in AMD (Yates et al., 2007), have been identified as successful drug targets for the disease (Kim et al., 2021).

As ClinVar and GWAS catalog provide such AMD-related genes, we used our Snp2Trait model to predict a list of SNPs that are potentially associated with AMD. Figure 3 (a) provides the SNPs and the genes involved that were predicted to be associated with AMD. The first three rows show that the model correctly confirms the top known associations of AMD with HTRA1, ARMS2, CFH, and CPN1. The final row of Figure 3 (a) highlights IL-27, a gene potentially associated with AMD but not reported in the GWAS Catalog or ClinVar. To dig deeper into the biomedical entity network, including AMD, *IL-27*, and *CPN1*, a known but under-explored gene associated with AMD (Han et al., 2020), we ran the literature evidence mining pipeline to collect relevant manuscripts and extract entities and their relations from the text.

**Figure 3.**
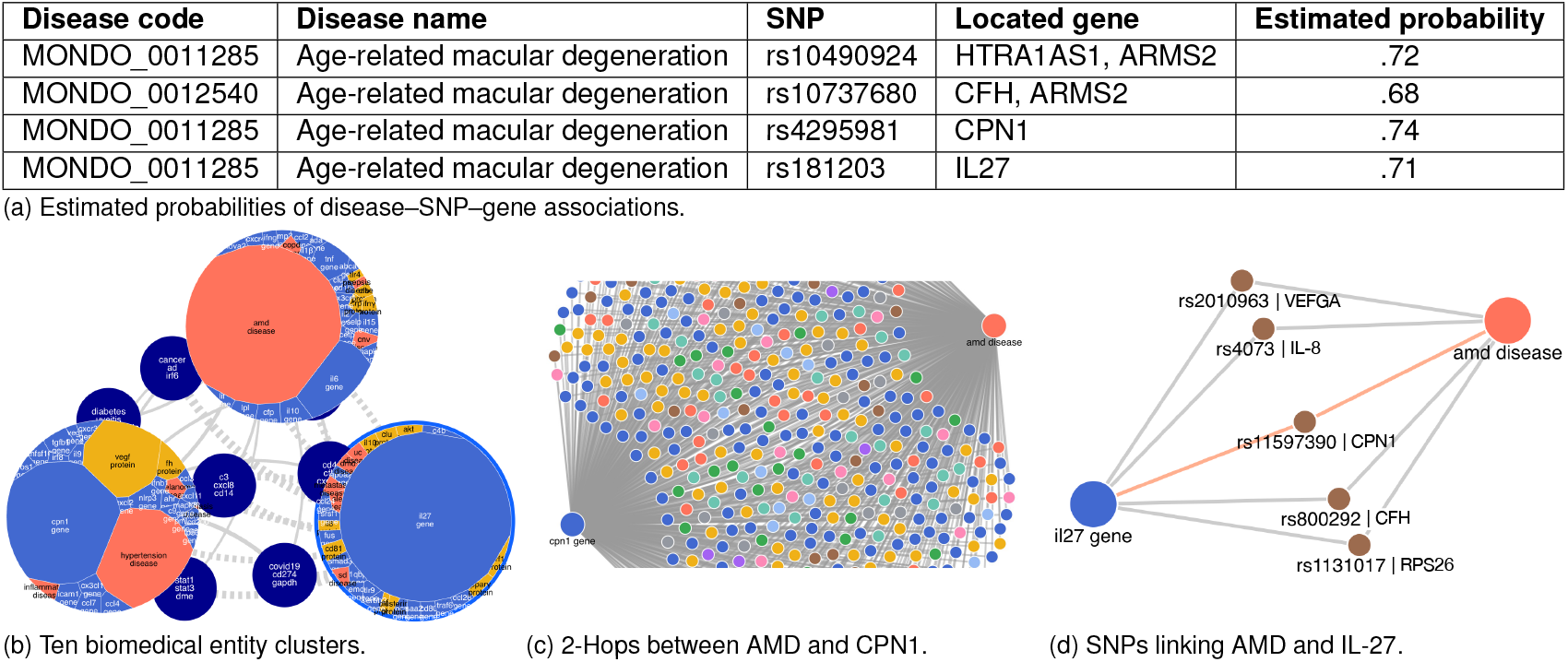
Description of the GENET workflow applied to AMD. (a) We first ran the Snp2Trait model to confirm associations between AMD and known SNPs/genes (top two rows) and then to identify hypothesized associations (bottom two rows). The estimated probabilities are in a similar range, which led us to launch the literature-mining pipeline for the three entities (AMD, IL-27, CPN1) separately and combine them. (b) With the combined knowledge graphs, we launched the visualization, which shows 10 clusters in total. Three clusters are expanded to inspect the entities that coexist with each of the three target entities within their respective clusters. The 2-hop connection view between AMD and CPN1 shows a variety of entities between them. (d) In the focused view between AMD and IL-27, we identified five SNPs and genes connecting AMD and IL-27. Along with well-known genes such as CFH and VEFGA, we found that CPN1 also bridges between AMD and IL-27.

To support our hypothesis and find direct or indirect connections between AMD, CPN1, and IL27, we ran three independent runs of literature mining and entity-relation extractions using each of these three keywords on PubMed Central. As a result, we have identified 28,400 papers mentioning AMD, 173 mentioning CPN1, and 5,284 papers mentioning IL-27. Subsequently, for this set of manuscripts, we extracted biomedical entities and relations, where each relation consists of one of the three keywords *AMD, IL-27, and CPN1*, so that we limit our search to the entities of interest, creating three star-shaped entity-centric knowledge graphs (KGs) (Theodoropoulos et al., 2023). The entities were first identified using a combination of the Unified Medical Language System (UMLS) (Bodenreider, 2004) and dictionary annotators (Kang et al., 2021), and then extracted relations using our pipeline with metallama3.3-70b as a large language model (LLM). In total, we extracted 400,205, 44,591, and 1,376 relations for AMD, IL-27, and CPN1, respectively (see Figure 3 (b)).

The three sets of entity-centric KGs were then combined into a single set to allow the identification of direct (one-hop) and indirect (two-hop) connections between AMD, IL-27, and CPN1. In total, we discovered 33,857 papers in PubMed Central, where we extracted 114,693 unique entities and 446,172 relations among them. We did not find a direct connection between AMD and CPN1 gene or between IL-27 and CPN1 genes, while there were 14 direct relations between AMD and IL-27 genes. In addition, 2,989 intermediate entities connected AMD and IL-27 gene, 24 entities connected IL-27 and CPN1 genes, and 215 two-hop entities between AMD and CPN1 (see Figure 3 (c)).

We ran clustering with K-Means (N=10) to identify groups of entities and visualized the network using GENET, as Figure 3 (b) shows. The visualization provides an overview of the relation network among the three main biomedical entities and their expanded relations with many other entities. By collecting the evidence and exploring the direct and expanded relations among entities, reading the actual text that indicates such relations, and visualizing entity-specific information for SNPs, proteins, and molecules, again, we did not find direct evidence that CPN1 is implicated in AMD. Thus, we devise a mechanism for its involvement through other biomedical entities. Figure 3 (c) shows an overview of 2-hop relations between AMD and CPN1. Among many biomedical entities that are independently connected to AMD and CPN1, inflammation, for instance, is an entity that is connected to CPN1 as a potential regulator (Matthews et al., 2004) and plays an important role in the onset and progression of AMD (Arrigo et al., 2023).

In particular, as Figure 3 (d) shows that the literature search indeed finds that SNP *rs11597390* (Carboxypeptidase N Subunit 1: CPN1) connects IL-27 to AMD. Consequently, the model and literature search provide hypotheses that complement each other, as an expansion of the search to 2-hop relationships brought agreement between the SNP2trait model and the evidence pipeline regarding an established relationship between AMD and CPN1. The actual manuscript describing a relationship between CPN1 and AMD (Han et al., 2020) was not found because our literature search of full texts was restricted to Pubmed Central and not PubMed in general. Thus, GENET not only generates novel hypotheses but also triangulates them with both model predictions and literature search, enabling a systematic expansion of the investigative horizon to enhance the reliability of hypothesis formulation. Overall, we find that the mechanism by which IL-27 potentially affects AMD is through CPN1 inactivation of components C3a and C5a (Matthews et al., 2004), and as ligation of C5aR inhibits IL-27 secretion (Bosmann et al., 2012), which either directly impacts AMD disease or activates CFH (Yoshida and Hunter, 2015).

## Conclusion

GENET demonstrates that an AI-powered, low-code visual analytics workflow can streamline the end-to-end process of extracting and exploring biomedical entity networks hidden in scientific literature and databases and help discover potential new drug targets, such as IL-27, when we applied GENET to age-related macular degeneration. By integrating AI-driven biomedical network analysis, LLM-based literature mining, clustering, and interactive visualizations, GENET creates a seamless human-in-the-loop environment in which domain experts supply a disease or genomic target, in this case derived from a model, and then iteratively explore, validate, and refine hypotheses through intuitive visual interfaces. This design frees scientists from the labor-intensive tasks of data wrangling and workflow orchestration, allowing them to devote their expertise to interpreting entity networks, uncovering novel mechanistic pathways, and designing follow-up experiments.

## Notes

### Competing Interest Statement

The authors have declared no competing interest.

